# Boundary-Driven Oscillations Rescue PdsA^-^ cells

**DOI:** 10.1101/752014

**Authors:** T. Eckstein, E. Vidal-Henriquez, A. Gholami

**Author notes:** These authors contributed equally (T. Eckstein, E. Vidal-Henriquez).

## Abstract

*Dictyostelium discoideum* amoeba aggregate if deprived of nutrients, producing cAMP waves at precisely timed intervals. Degradation of extracellular cAMP by the enzyme phosphodiesterase PdsA is fundamental to successfully producing waves, regulating the external cAMP gradient field and preventing the accumulation of cAMP. The knockout mutant PdsA^-^ produces no or a greatly reduced amount of main extracellular phosphodiesterase, therefore failing to relay cAMP waves and aggregate under starvation conditions. Using a microfluidic channel, we show how an advective flow can partially recover signaling in a population of starving PdsA^-^ cells. Above a minimum flow velocity, decaying waves are induced, with a decay length that increases with the imposed flow velocity. Interestingly, after stopping the advecting flow, the cells continue to signal, showing wave propagation and aggregation, although with a wave period much higher than in wild type cells. We performed extensive numerical simulations and showed that these waves have a boundary-driven origin, where the lack of cAMP in the upstream flow destabilizes the system. We explored the properties of these waves and the parameter region where they exist, with good agreement with our experimental observations. These boundary-driven waves dominate the system dynamics in the velocity range where they exist, while at higher flow velocities the natural wave period of 6 min recovers. These results provide experimental confirmation of the destabilizing effect of the upstream boundary in an otherwise stable reaction-diffusion system. We expect this mechanism to be relevant for wave creation in other oscillatory or excitable systems that are incapable of normal pattern formation.

**SIGNIFICANCE STATEMENT:** We present experimental evidence for the existence of boundary-driven instabilities in a reaction-diffusion-advection system. In our theoretical prediction (1), we have shown that imposing an absorbing boundary condition on the upstream end of a flow-through channel filled with signaling cells creates an instability capable of periodically producing wave trains which are advected downstream. Under starvation, these cells secret the signaling molecule cAMP as well as the degrading agent phosphodiestrase that degrades cAMP. This instability was predicted to exist at lower degradation rates of cAMP and thus was expected to provide a mechanism for wave creation in phosphodiesterase deficient systems, such as PdsA^-^ cells. Our experiments confirm the importance of the upstream boundary condition and show that boundary-driven oscillations are relevant in reaction-diffusion systems.

## INTRODUCTION

The social amoeba *Dictyostelium discoideum (D.discoideum)* is a paradigm model organism to study biological pattern formation and provides insights into the nature of cell-cell communication and emergent collective behavior (2–5). *D. discoideum* cells lead a solitary life as long as the food supply is sufficient. Upon nutrient depletion the cells start an aggregation process in which they gather in groups of around 10^4^ – 10^6^ amoebas and form multi-cellular structures known as fruiting bodies in order to survive (6, 7). This aggregation process is governed by the chemotactic response of *D. discoideum* cells to outward propagating cAMP (cyclic adenosine monophosphate) waves that trigger oriented periodic movement of the cells towards the aggregation centers. These waves are generated by an interplay between random diffusion of cAMP in the extracellular medium and the ability of the cells both to receive the signal via cAMP receptors on the membrane of individual cells as well as to relay and amplify the signal by producing more cAMP. The cells crawl up the concentration gradient for a few minutes until the chemotactic response of the cells is adapted (8).

The limiting factor in signal amplification is the desensitization of cAMP receptors if they are persistently exposed to high concentrations of extracellular cAMP (9–11). To avoid receptor desensitization and to produce strong cAMP gradients, the cells emit phosphodiesterases (PDEs) that degrade the extracellular cAMP. Three extracellular PDEs have been characterized (12) in *D. discoideum*: PDE1 (also called PdsA or PdeA), PDE4, and PDE7. All three types of PDEs have different dynamics, therefore becoming more relevant for cAMP degradation during different parts of the developmental program. During the early aggregation stage, PdsA is the dominant PDE, degrading the extracellular cAMP almost all by itself. The knockout mutant PdsA^-^ does not produce PdsA and has been shown to be unable to produce cAMP waves, and thus fail to aggregate (13). Rescue of this cell type was reported by adding external phosphodiesterase from a rat brain (14), PDE overexpressors (13) or by mixing with WT cells (15). Moreover, it has been shown that PdsA^-^ amoebas show oscillations in the concentration of intracellular cAMP if subjected to a fresh buffer flow that carries extracellular cAMP away (16), suggesting that the external flow can replace the role of PDE.

In their natural habitat, *D. discoideum* cells are exposed to external flows which can significantly influence the wave generation process (17). Previously, we had numerically predicted the existence of a convective instability induced by the influx of cAMP-free buffer through a colony of signaling amoebas (1, 18). In such a system it is of utmost importance what kind of chemicals get injected in the system with the advecting flow. In the numerical simulations this is equivalent to what boundary conditions are used in the upstream edge of the system. In the case where there are no cells upstream, the flow would be free of cAMP and can act as a destabilizing agent. We have shown that holding the upstream boundary to a zero concentration of cAMP produces an instability that periodically sends wave trains downstream. The wave generation mechanism works by first advecting the cAMP downstream, thus depleting the upstream area of cAMP. This low concentration of the chemoattractant destabilizes the cells close to the upstream boundary which react by releasing a pulse of cAMP. This instability, known as boundary-driven oscillations (1), exists at lower degradation rates than the oscillatory regime and thus provides a mechanism for wave creation in phosphodiesterase-deficient systems, such as PdsA^-^ cells.

In this work, we present experimental evidence of the existence of boundary-driven oscillations (BDOs), which can be observed in a microfluidic setup filled with starving PdsA^-^ cells. Note that the imposed flows are not strong enough to detach the cells from the substrate and the cAMP produced by the cells is advected downstream. Interestingly, at small flow velocities we observe decaying cAMP waves that do not fill the whole length of the channel, with a decay length that grows with the advecting flow velocity. Our extensive numerical simulations confirm a similar trend in the decay length of the waves as the flow is increased. We also quantified the period of these boundary-driven waves, measuring high wave periods (~ 25 min) at small advective flows which decreases with the imposed flow velocity, approaching the natural period of 6 min at flow velocities larger than 0.6 mm/min. Interestingly, once the flow is switched off, the cells located upstream of the microfluidic channel (that have experienced BDOs) continue to produce cAMP waves similar to wild type cells, but with a much larger wave period of 15-25 min. These amoebas are able to aggregate normally, thus the aggregation phenotype is rescued. However, cells downstream of the channel located outside the penetration depth of BDOs fail to aggregate once the flow is switched off. Thereby, at flow velocities larger than 1 mm/min where the BDOs fill the whole length of the channel, all the cells in the channel are rescued. We observed a similar phenomenon as we used WT cells at the upstream area of the channel. The cAMP waves generated by WT cells penetrate gradually throughout the population of PdsA^-^ cells, covering the whole length of the microfluidic channel.

## MATERIALS AND METHODS

### Cell Culture

All experiments were performed with *D. discoideum* UK5 cells, denoted as PdsA^-^ (Dicty Stock Center, Chicago, IL, USA). Cells were grown in HL-5 medium (35.5 g of Formedium powder from Formedium Ltd, Norfolk, England, per liter of double-distilled water, autoclaved and filtered) at 22 °C on polystyrene Petri dishes (TC Dish 100, Sarsted, Nümbrecht Germany) and harvested when they became confluent. Before the experiments, the cells were centrifuged and washed two times with phosphate buffer (2 g of KH_2_PO_4_ and 0.36 g of Na_2_HPO_4_.H_2_O per liter at pH 6.0, autoclaved, both from Merck, Darmstadt, Germany). The cells were then resuspended in 200 *μ*l phosphate buffer. The cell density was determined using a hemocytometer (Neubauer Zählkammer, VWR, Darmstadt, Germany), diluted to 5 × 10^7^ cells/ml of phosphate buffer and injected into the microfluidic channel.

### Microfluidics

The microfluidic devices were fabricated by means of standard soft lithography methods (19). A silicon wafer was coated with a 100 *μ*m photoresist layer (SU-8 100, Micro Resist Technology GmbH, Berlin, Germany) and patterned by photolithography to obtain a structured master wafer. The channels were 2 mm wide, 50 mm long, and 103 ± 2 *μ*m high. Polydimethylsiloxane (PDMS, 10:1 mixture with curing agent, Sylgard 184, Dow Corning GmbH, Wiesbaden, Germany) was poured onto the wafer and cured for 1 h at 75°C. To produce the microfluidic device, a PDMS block containing the macro-channels was cut out, and two inlets (7 mm and 0.75 mm in diameter) were punched through the PDMS at opposite ends of the channel with the help of PDMS punchers (Harris Uni-Core-7.00 and Harris Uni-Core-0.75, Sigma-Aldrich, St. Louis, MO, USA). Afterwards, a glass microscope slide (76×26 mm, VWR, Darmstadt, Germany) was sealed to the PDMS block following a 20–30 s treatment in air plasma (PDC 002, Harrick Plasma, Ithaca, USA) to close the macro-channels (see supplementary figure S1). The large inlet (7 mm diameter) was used as a liquid reservoir filled with phosphate buffer and from the other side phosphate buffer was sucked out using a high precision syringe pump (PHD 2000 Infuse/Withdraw Syringe Pump from Harvard Apparatus, Holliston, USA, combined with gas-tight glass syringes from Hamilton, Reno, USA) at constant buffer flow rates.

### Image Acquisition and Analysis

We used a dark-field setup consisting of a monochrome 12-bit CCD camera (QIClick-F-M-12, QImaging, Surrey, Canada), a telecentric lens (0.16X SilverTL Telecentric Lens, Edmund Optics Inc, Barrington, NJ, USA) and a ring light source (Advanced Illumination, Rochester, VT, USA). The camera was controlled with an image capture program (Micro-Manager (20)) and it recorded images every 20 seconds. We processed the dark-field videos by first subtracting the intensity trend of each pixel to normalize the signal in time. Next, we did spatial band-pass filtering to suppress structures bigger than 3.5 mm and smaller than 0.294 mm. Finally, we used the Hilbert transform (21) to calculate the phase of each pixel, thus obtaining the phase maps.

### Numerical Simulations

The Martiel-Goldbeter model (22) is a reaction-diffusion model proposed in 1987 to describe pattern formation in *D. discoideum* and has been successfully used to describe spiral formation (3,23), flow-driven waves (24), spontaneous formation of aggregation centers (25), and cell streaming (26). Changing the parameters in the model allows us to describe the system at different points during the starvation process or with knocked-out genes. To describe PdsA^-^ cells, then, we use this model with a very low cAMP degradation rate.

Numerical simulations of the Martiel-Goldbeter model (22) to describe cAMP production and relay in *D. discoideum* were performed and compared with experimental results. The relay process starts when the cells detect the presence of cAMP in the extracellular medium through the receptors located on the cell membrane. After binding with cAMP, the receptors change to a phosphorylated or inactive state, in which they have a lower probability of binding with cAMP. After some relaxation time, the receptors go back to their active state. The binding of the receptors with cAMP triggers a series of reactions inside the cell that culminate with the production of cAMP and the cells’ motion against the wave direction. The produced cAMP is degraded into AMP by the intracellular phosphodiesterase and transported passively to the extracellular medium. Finally, once outside of the cells, the cAMP is free to diffuse and is degraded by the action of phosphodiesterase in the extracellular medium. This is the type of phosphodiesterase that PdsA^-^ cells do not produce.

The equations of the model are

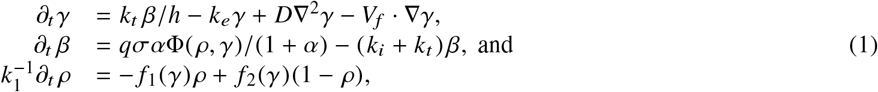

where the modeled fields are the extracellular concentration of cAMP *γ*(*x, t*), the intracellular concentration of cAMP *β*(*x, t*), and the percentage of active receptors on the cell membrane *ρ*(*x, t*). *D* stands for the cAMP diffusion, *V_f_* the velocity of the externally applied flow, Φ is the nonlinear function for cAMP production, *f*_1_ the function for receptors phosphoryliation and *f*_2_ for receptors resensitization, these functions are given by

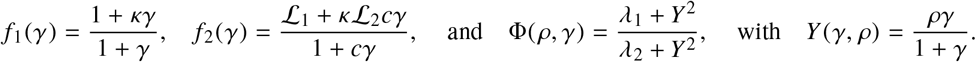

All used parameters are indicated in Table 1, and were kept fixed with the exception of *σ* (linearly proportional to the production rate) and *k_e_* (degradation rate) which were used to explore the parameter space. To reproduce the signaling process in PdsA^-^ cells, *k_e_* was kept low, *k_e_* ≪ 1 min^-1^, compared to simulations of wild type cells where *k_e_* ≈ 5 – 12 min ^-1^(3, 22, 23).

**Table 1:** Parameters used in numerical simulations of Equation 1

|  |  |  |
| --- | --- | --- |
| $c = 10$ | $h = 5$ | $k_1 = 0.09 \text{ min}^{-1}$ |
| $\kappa = 18.5$ | $\alpha = 3$ | $k_i = 1.7 \text{ min}^{-1}$ |
| $k_t = 0.9 \text{ min}^{-1}$ | $\mathcal{L}_1 = 10$ | $\mathcal{L}_2 = 0.005$ |
| $q = 4000$ | $\lambda_1 = 10^{-4}$ | $\lambda_2 = 0.2575$ |
| | $D = 0.024 \text{ mm}^2/\text{min}$ | |

Numerical simulations were performed using finite differences with a 3 points Laplacian in 1-D and 5 points in 2-D for the space discretization and a Runge-Kutta scheme with an adaptative time step (27) for the time evolution. A nonlinear discretization of the advection operator was used to ensure positivity of the cAMP concentration *γ* following the work of Koren (28). To account for the injection of buffer without cAMP an absorbing boundary condition was used, that is, the value of *γ* was fixed at zero at the upstream boundary, *γ*(*x* = 0, *t*) = 0. At the downstream boundary no-flux (*∂_x_γ*(*x* = *L, t*) = 0) boundary conditions were used. For a detailed analysis of the effects of advection in this configuration with WT cells, refer to our previous works (1, 24).

## RESULTS

### A fresh cAMP-free buffer flow can substitute PdsA^-^ activity

In a population of starving *D. discoideum* cells, the level of extracellular cAMP is controlled by the activity of the enzyme adenylyl cyclase (ACA), which catalyzes the reaction in which cAMP is produced, and phosphodiestrase, which degrades cAMP. It is believed that PdsA, by degrading extracellular cAMP, plays an important role in regulating chemotaxis by establishing cAMP gradients during the streaming process. Upon starvation, PdsA^-^ cells fail to produce cAMP waves and thus can not aggregate. However, in our experiments with flow-through microfluidic channels, we observe that an external flow can effectively play the role of PdsA by removing the extracellular cAMP. Thereby, cAMP waves are recovered and the aggregation process is rescued.

We observe boundary-driven waves in a population of PdsA^-^ cells if they experience the external flow for at least three hours. The minimum imposed flow velocity needed to recover cAMP waves in our experiments was *V_f_* ≥ 0.3 mm/min. At velocities below 0.3 mm/min we did not observe any waves even during our long-run experiments where buffer was flowing through the channel for more than 10 hours. Figure 1a-c shows examples of boundary-driven waves at low, moderate and high imposed flow velocities, respectively. In all of our experiments, the cAMP waves develop within three hours of imposing the flow. While the waves at *V_f_* = 0.3 mm/min decay quickly as they initiate at the upstream end of the channel and never successfully propagate throughout the channel, waves at *V_f_* = 0.7 mm/min eventually are able to penetrate and travel along the whole length of the channel. Full penetration of the waves in the channel at moderate flow speeds, occurs within the time scale of three hours and the transition from partial to full penetration is fairly sharp. Interestingly, at high flow speed of *V_f_* = 1.5 mm/min, once the waves develop upstream, they are able to propagate downstream throughout the channel almost immediately. We systematically measured the decay length of the waves as we increased the imposed flow velocity and found that it grows with the flow. Figure 2a-d summarizes properties of these boundary-driven waves in terms of their wave velocity, decay length, period, and wavelength. Below, we consider different flow regimes and discuss wave properties in detail.

**Figure 1:**
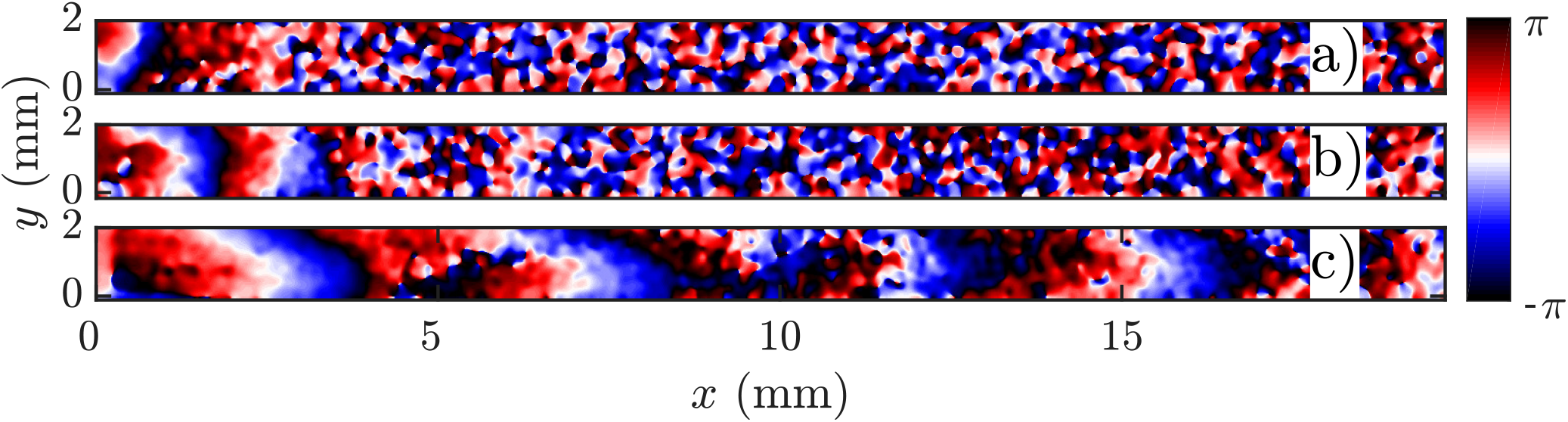
Phase map of boundary-driven waves in a population of PdsA^-^ cells at a) low flow speed of *V_f_* = 0.3 mm/min, b) moderate flow speed of *V_f_* = 0.7 mm/min and c) high flow speed of *V_f_* = 1.5 mm/min. While the waves in a) and b) are almost planar and decay along the channel, the waves in c) are parabolic and fill the whole length of the channel (see supplementary movies S1-S3). All images are shown 300 min after the flow was switched on at *t* = 0.

**Figure 2:**
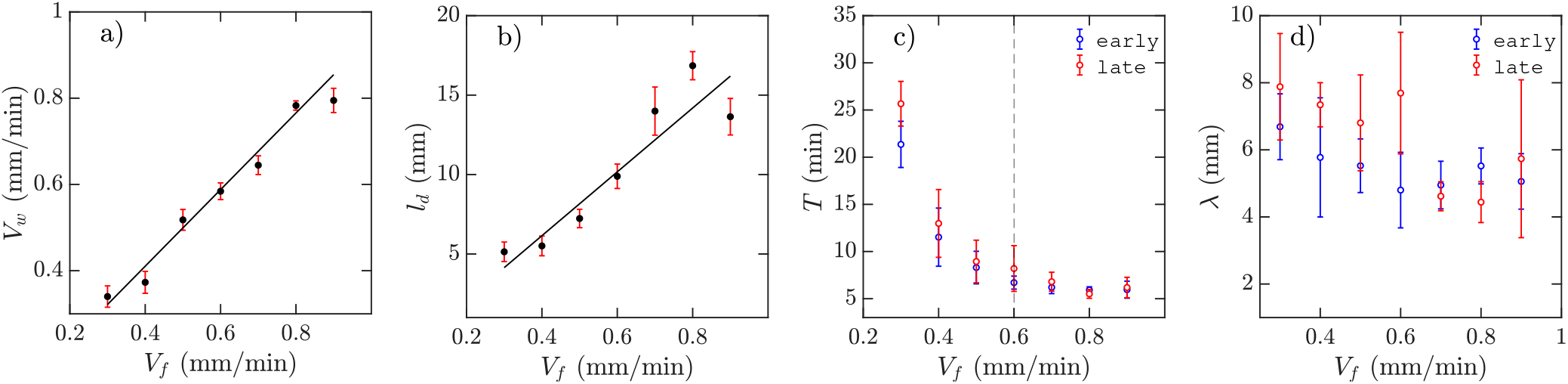
Experimental characterization of boundary-driven waves relayed by PdsA^-^ cells when subjected to a flow of cAMP-free buffer. a) Wave velocity *V_w_*, b) wave decay length *l_d_*, c) wave period *T*, and d) the wavelength *λ* as a function of the imposed flow velocity *V_f_*. Solid black lines in a-b) represent a linear least-squares fit to the data. Blue and red data points in c-d) are from the early and late regime of the experiments. For each data point, minimum three experiments are included to calculate the mean and the standard deviation.

### A. Waves at low flow velocities

The development of boundary-driven waves at the low flow velocity of *V_f_* = 0.3 is shown in Fig. 3 (see supplementary movie S1). A wave front initiates at the upstream boundary of the channel and decays quickly over a length scale of ~5 mm. Over time, the waves successfully travel a longer distance (~ 10 mm) before dying out. We define a characteristic length *l_d_* which corresponds to the length that a wave travels inside the channel before its amplitude is lower than 10% of its maximum amplitude. This length scale grows with the applied flow speed, meaning that the waves reach further downstream in the channel at higher flow speeds. Based on the decay length of the waves at a given flow velocity, we define an early and a late regime. During the early regime, the waves decay after a shorter length than during the later regime. Therefore, we systematically measure the wave period and the wavelength corresponding to each regime separately, as shown in Fig. 2c-d.

**Figure 3:**
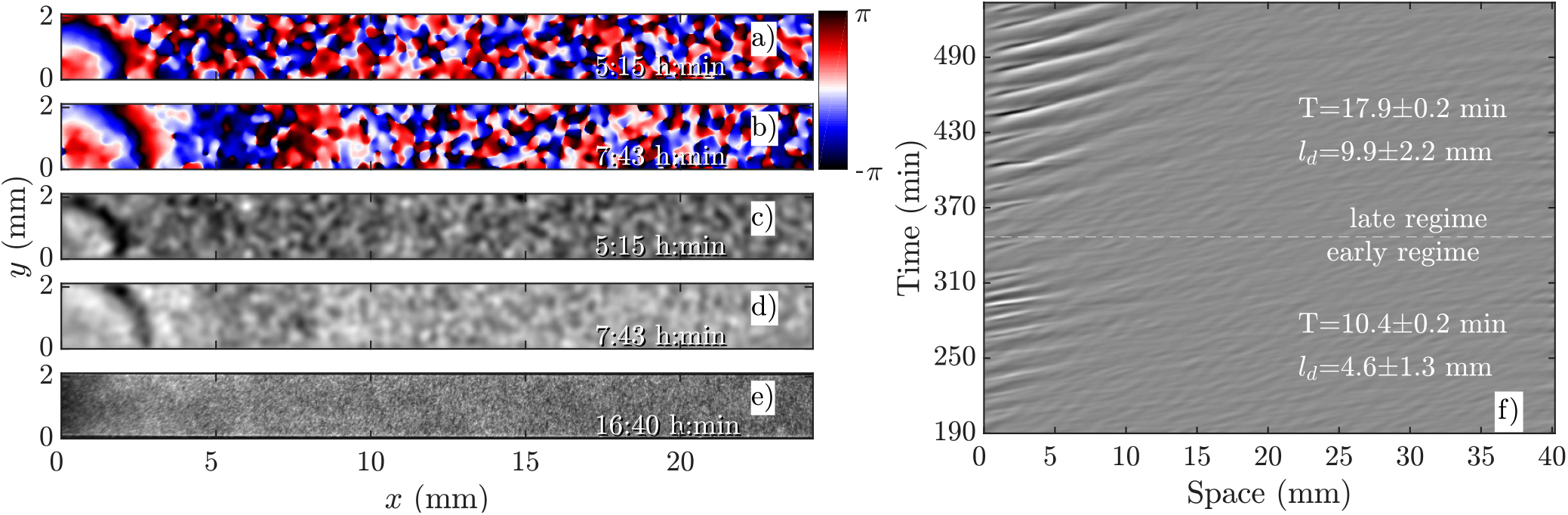
Overview of an experiment at a low flow speed of *V_f_* = 0.3 mm/min (see movie S1). a-b) phase images extracted from processed images in c-d). a) and c) show waves during the early regime that decay immediately upon entering the channel. b) and d) show waves during the late regime, which move slightly farther along the channel before decaying. e) Cells fail to aggregate even at the end of the experiment. f) shows part of the space-time plot of this experiment. Timestamps denote time since the flow was switched on at *t* = 0.

An interesting feature of the waves developed in the flow range of 0.3 ≤ *V_f_* ≤ 0.6 is their large period. We observed periods as large as 20-25 mins for *V_f_* = 0.3 mm/min, decreasing to 10-15 min at *V_f_* = 0.4 mm/min and approaching the normal period of 6 min at *V_f_* = 0.6 mm/min (see Fig. 2c). It is plausible that at flow velocities larger than 0.6 mm/min, the period of the waves is set by the random firing of the cells and not anymore by BDOs. This hypothesis was confirmed by our numerical simulations. Finally, the aggregation pattern of the cells in the presence of boundary-driven waves for *V_f_* = 0.3 mm/min is shown in Fig. 3e. It seems that the cells even at the upstream part of the channel that were exposed to boundary-driven waves fail to aggregate. We believe that this is due to the fact that the amplitude of the boundary-driven waves at low flow speeds is not strong enough to trigger aggregation of the cells. The minimum flow speed needed in our experiments to rescue aggregation of the cells was 0.6 mm/min. Note that this velocity coincides with the velocity at which period of boundary-driven waves sets to the natural period of 6 min (see the dashed line in Fig. 2c).

### B. Waves at moderate flow velocities

As we increase the flow velocity, in the range 0.6 mm/min≤ *V_f_* ≤ 1 mm/min, waves behave differently. Although in the early phase, they propagate only in part of the channel (Fig. 4a,c), in the late phase they successfully propagate throughout the channel (Fig. 4b,d). Moreover, the waves are stronger in amplitude and have the normal period of 6 min, similar to the period of the flow-driven waves in WT cells (17, 24). Regarding the aggregation pattern, we observe clusters of cells mostly in the upper part of the channel, confirming that aggregation is partially rescued. We also performed bright-field measurements to look closely at aggregation of PdsA^-^ cells in the presence of flow (*V_f_* = 1 mm/min). Figure 5 shows that rescue of the cells is more successful at the upstream end of the channel where the boundary-driven waves penetrate initially and have a larger amplitude (see supplementary movie S4). Further downstream, the amplitude of the waves decay and is not strong enough to rescue the cells. At higher flow speed, aggregation clusters cover a larger portion of the channel. We emphasize that cells do not show aggregation clusters if the flow is stopped before they start producing boundary-driven waves.

**Figure 4:**
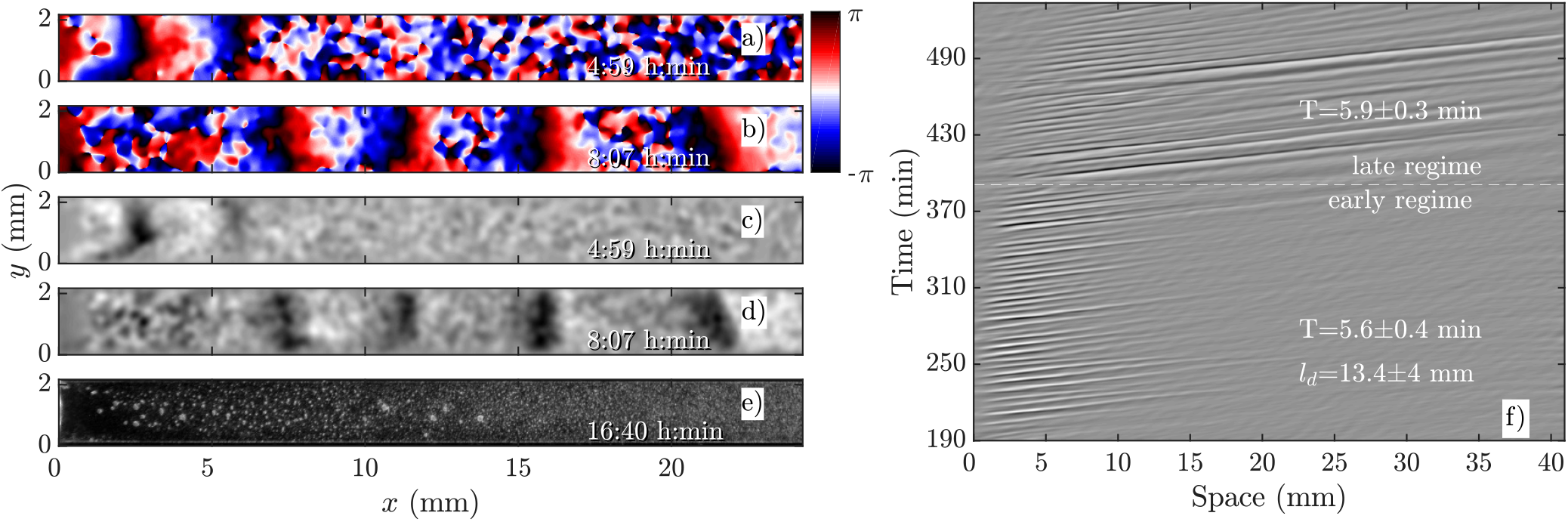
Overview of an experiment at a medium flow speed of *V_f_* = 0.7 mm/min (see movie S2). a-b) phase images extracted from processed images in c-d). a) and c) show waves during the early regime that decay quickly along the channel. b) and d) show waves during the late regime, which fill the entire channel. e) shows aggregation at the end of the experiment. f) shows a part of the space-time plot of this experiment. Timestamps denote time since the flow was switched on at *t* = 0.

**Figure 5:**
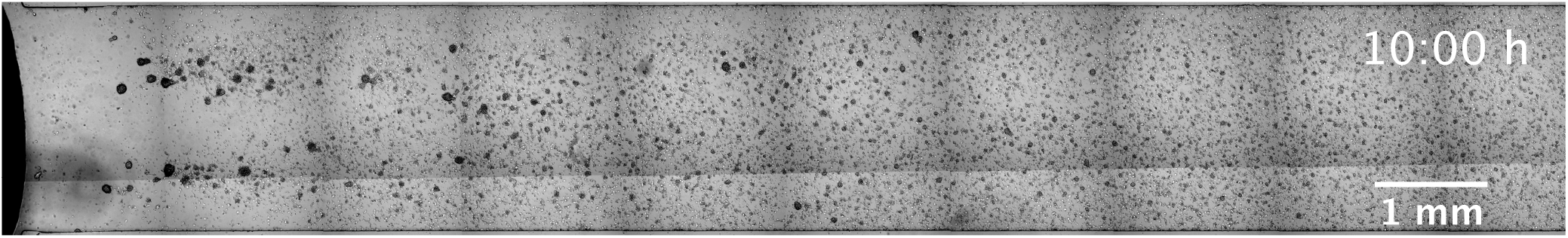
A bright-field image showing aggregation of cells in a flow-through channel. Images are taken at 4x magnification at different parts of the channel and stitched together. Note that at the upstream part of the channel we have a reservoir filled with phosphate buffer which is free of the cells and cAMP (see movie S4).

To examine the effect of different imposed flow velocities in a single experiment, we constructed a channel with a sidearm as shown in Fig. 6. Since flow resistance in the sidearm is higher than the direct path, the flow in the sidearm is about 2.2 times slower. This means that a direct comparison of the behavior of flow-driven waves at different flow speeds is possible in one experiment. As expected, the waves in the direct channel penetrate longer and are stronger than the ones in the sidearm, as shown in Fig. 6a and supplementary movie S5. Similar to flow-driven waves in WT cells (24), the width of the wave fronts is larger at the straight part of the channel in comparison to the waves in the sidearm. Finally, the spatial extent of the waves controls the aggregation efficiency in the two channel parts. While the cells in the main channel aggregate quite well, there is a little aggregation in the sidearm where the spatial extension of the waves is shorter (see Fig. 6b). Note that in both parts of the channel, further downstream the aggregation process appears weaker compared to upstream, which we believe is related to the wave amplitude that decays as it propagates along the channel.

**Figure 6:**
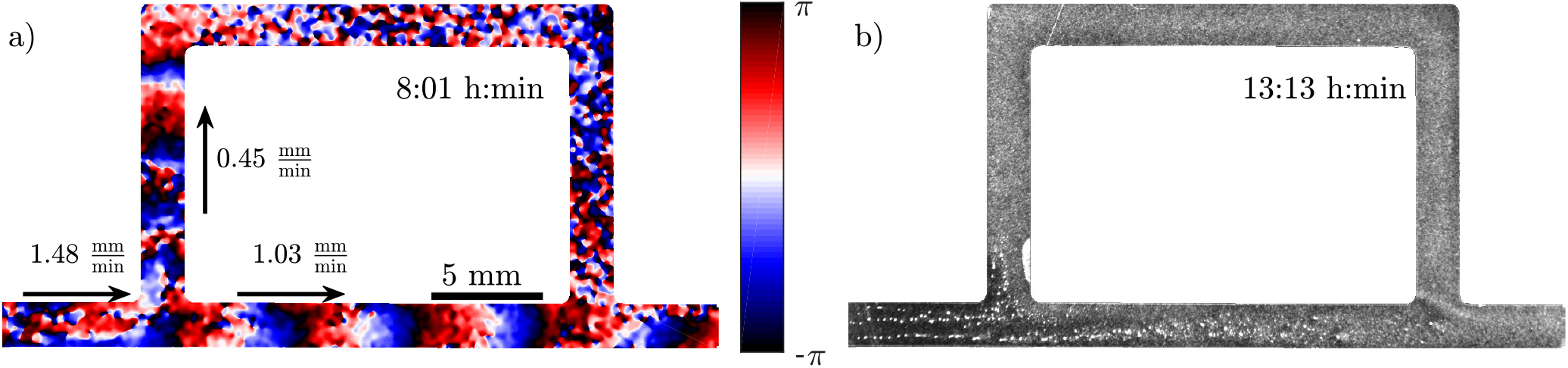
Experiment in a channel with a sidearm which has a lower flow speed than the straight part of the channel (see movie S5). a) Phase map of the developed boundary-driven waves and b) the corresponding aggregation pattern at the later times. Note different spatial extension of the waves in both parts of the channel which is the important factor in deciding if the aggregation phenotype will be rescued or not.

### C. Waves at high flow velocities

If the imposed flow velocity is increased even further, *V_f_* ≥ 1 mm/min, the so-called early phase disappears. This means that once the boundary-driven waves initiate at the upstream part, they propagate through the entire length of the channel. Figure 7 shows an experiment at *V_f_* = 1.5 mm/min. The waves have the normal period of 6 min and wave fronts are more elongated in comparison to small flow experiments, which is similar to flow-driven waves at high speeds described previously for WT cells (24). Cells over the entire length of the channel are rescued and successfully aggregate (see Fig. 7e and supplementary movie S3).

**Figure 7:**
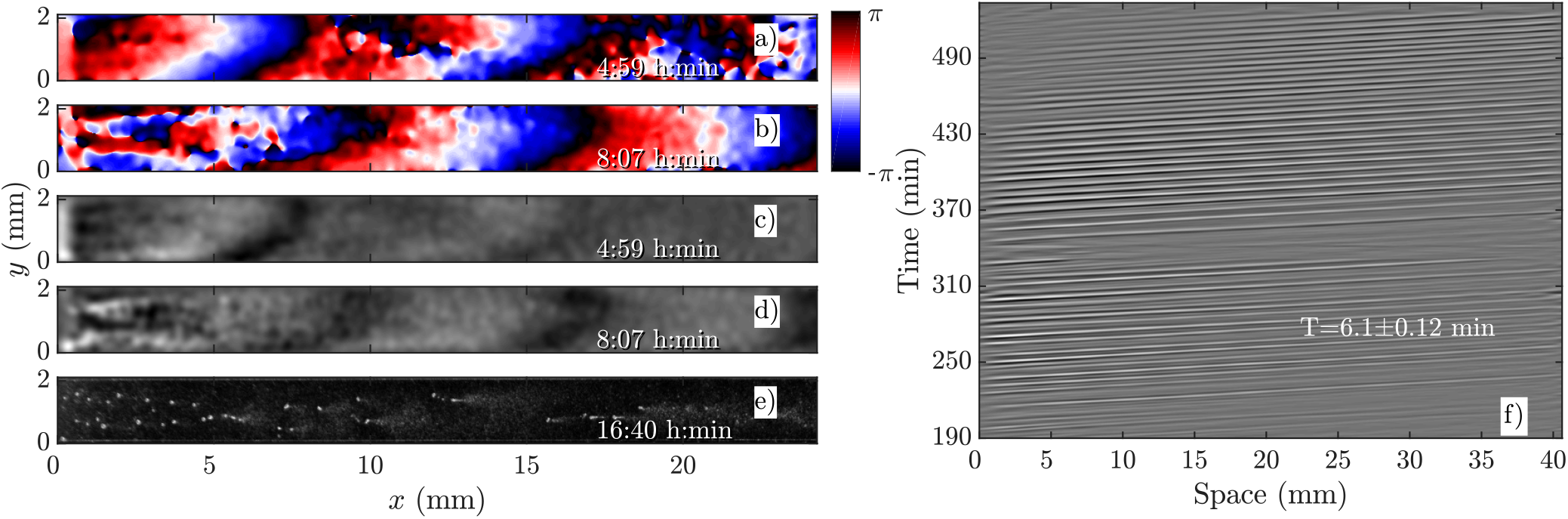
Boundary-driven waves at *V_f_* = 1.5 mm/min (see movie S3). a-b) Phase maps extracted from processed images in c-d). e) Cells over the entire length of the channel are rescued and aggregate normally. f) Space-time plot of the waves showing that once the waves develop after three hours of flow, they travel through the entire channel. Timestamps denote time since flow was switched on at *t* = 0.

### Rescue of PdsA^-^ cells by flow

Another interesting observation in our flow experiments with PdsA^-^ cells was the rescue of pattern formation once the flow was switched off. We exposed the cells to a minimum flow velocity of *V_f_* = 0.6 mm/min and let the boundary-driven waves develop and propagate for a minimum time of one hour before we switch off the flow. Interestingly, we observed that in less than 90 mins wave centers developed spontaneously and relayed through the population. This occurred again at the upstream parts of the channel that were exposed to boundary-driven waves (see Fig. 8 and supplementary movie S6). The period of the spontaneous waves was significantly higher than the ones produced by wild type cells (16-24 min compared to 4-6 min in wild type cells). The wave velocity was measured to be 0.263 ± 0.006 mm/min, which is comparable to the wave speed of 0.4 mm/min in wild type cells. A systematic measurement of wavelength, period and propagation velocity of the spontaneous waves is presented in Fig. 8b-c. In a single experiment, we measure a wide range of wave periods varying from 15 min (as they appear) and increasing to 25 min at later times. This is shown in Fig. 8b.

**Figure 8:**
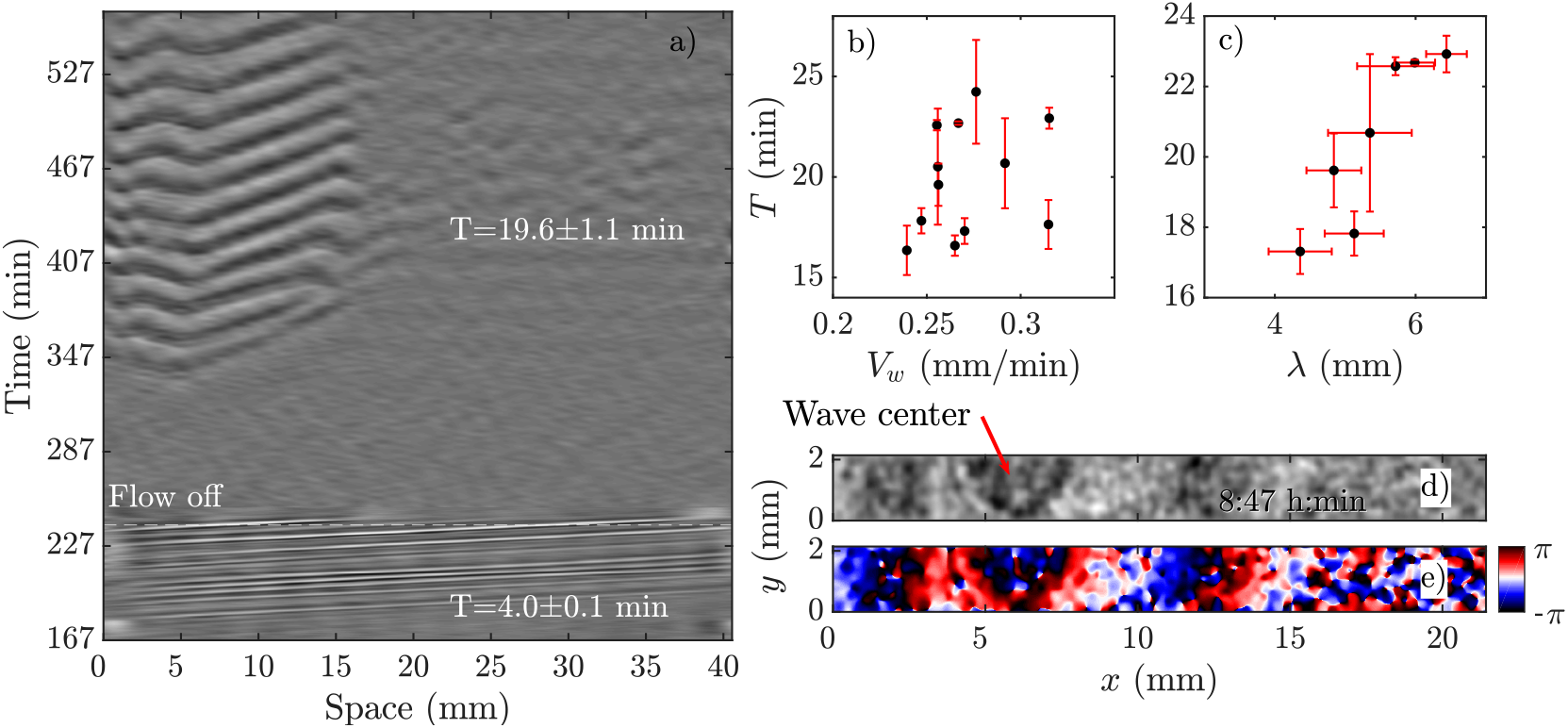
Pattern formation of PdsA^-^ cells after the flow was switched off. a) Space-time plot of an experiment at imposed flow of *V_f_* = 2 mm/min. The flow was turned off after 240 min (see movie S6). b) Period *T* and c) wavelength *λ* of spontaneous waves (appeared in the flow-off regime) plotted as a function of wave velocity *V_w_*. For these measurements set of different experiments at various flow velocities are used where the flow was switched off at different time points. d) Processed dark-field image and e) the corresponding phase map showing the initiation of a wave at the upstream part of a channel after the flow of *V_f_* = 2 mm/min has been switched off. The corresponding space-time plot of this experiment is shown in part a). Timestamp shows the time since the flow was switched on at *t* = 0.

Based on these results, we explored the minimum time PdsA^-^ cells need to be subjected to flow, such that they show spontaneous pattern formation once the flow is switched off. We found that the cells show spontaneous pattern formation, even if the population experienced the passage of very few boundary-driven waves before the flow was switched off. Figure 9a shows an example where the flow of *V_f_* = 2 mm/min was turned off once 1-2 pulses of boundary-driven waves pass through the channel. The cAMP waves form spontaneously at *t* ~ 360 min only at the upstream part of the channel. Interestingly, at higher flow speed of *V_f_* = 5 mm/min where we allow for boundary-driven waves to persist for about 2 hours, once we switch off the flow, the spontaneous waves appear quickly again at *t* ~ 360 mins. Since the flow-driven waves penetrate throughout the channel, the recovered spontaneous waves are also observed over the entire length of the channel (see movies S7,S8).

**Figure 9:**
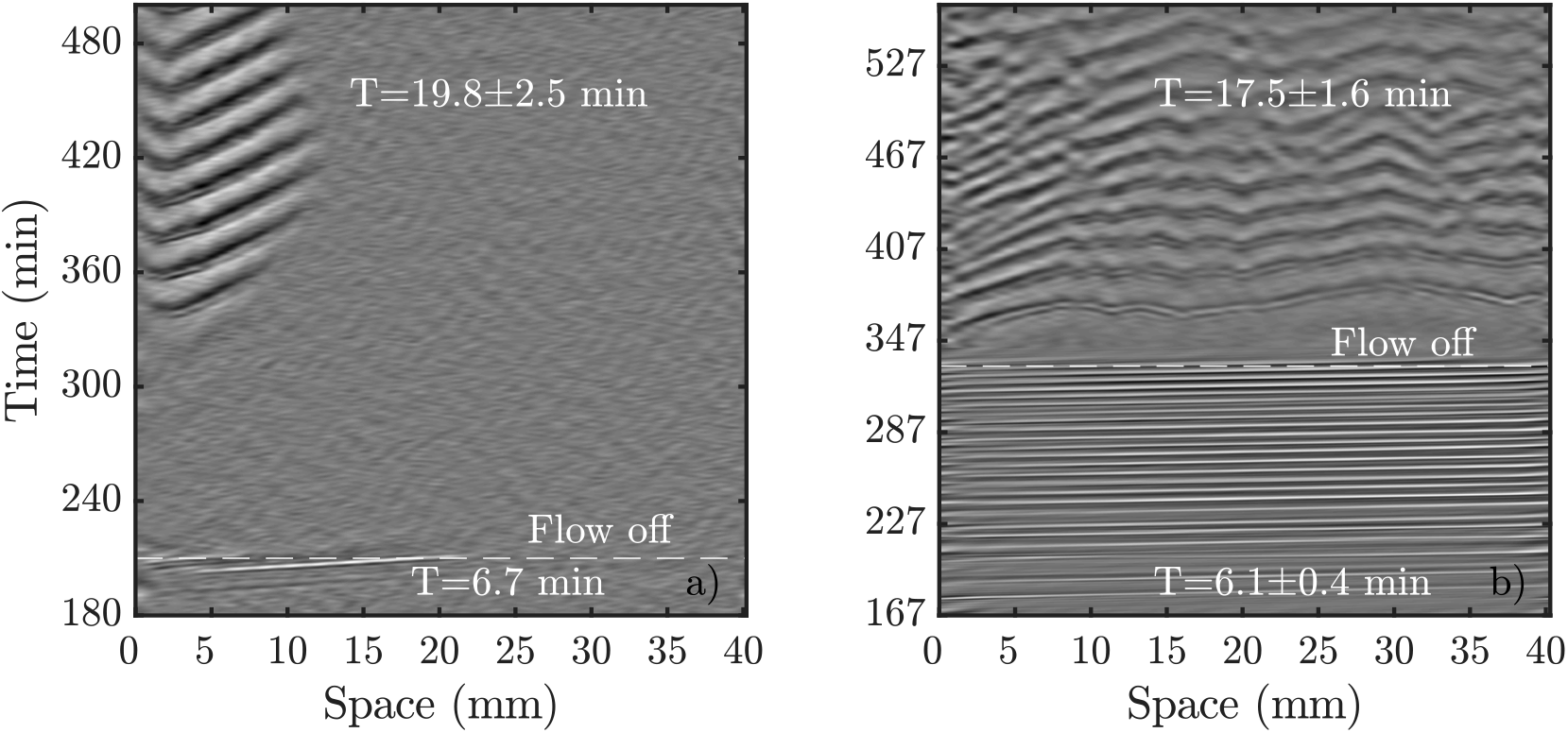
Comparison of spontaneous pattern formation once the flow is switched off. a) Flow of *V_f_* = 2 mm/min was switched off shortly after boundary-driven waves were observed. cAMP waves spontaneously appear at 360 min (see movie S7). b) Higher flow velocity of *V_f_* = 5 mm/min triggered boundary-driven waves that propagated through the population for more than two hours before the flow was turned off. The spontaneous waves appeared shortly after (see movie S8).

### Rescue of PdsA^-^ cells with Wild Type cells

As we mentioned above, pattern formation in PdsA^-^ cells is impaired because of the lack of cAMP degradation. Therefore, we tried rescuing the cells by using the PdsA produced by Ax2 cells. To do this, we injected PdsA^-^ cells into a channel and filled the reservoir of the channel with Ax2 cells. This reservoir is located upstream from the channel, and is where the fluid accumulates before flowing through the channel. Thus, the cell types were not mixed, but the buffer that flowed through the channel was preconditioned by its exposure to the Ax2 cells. We find that the behavior of this system depends strongly on the initial starvation time of the Ax2 cells used. In the case of using Ax2 cells without previous starvation, there is again no pattern formation for the first three hours of flow. Nevertheless, when waves emerge they do not decay along the channel, but instead fill the channel immediately for all flow speeds studied. However, if we use Ax2 cells that were initially starved for 4 hours, we find that pattern formation is observed immediately after the flow is switched on. Initially, the waves decay rather quickly, but the decay length increases exponentially over time until the whole channel is filled, see Fig. 10a and supplementary movie S9. The rate at which the decay length increases depends on the imposed flow velocity, filling the whole channel more quickly for higher advecting flows (see Fig. 10b).

**Figure 10:**
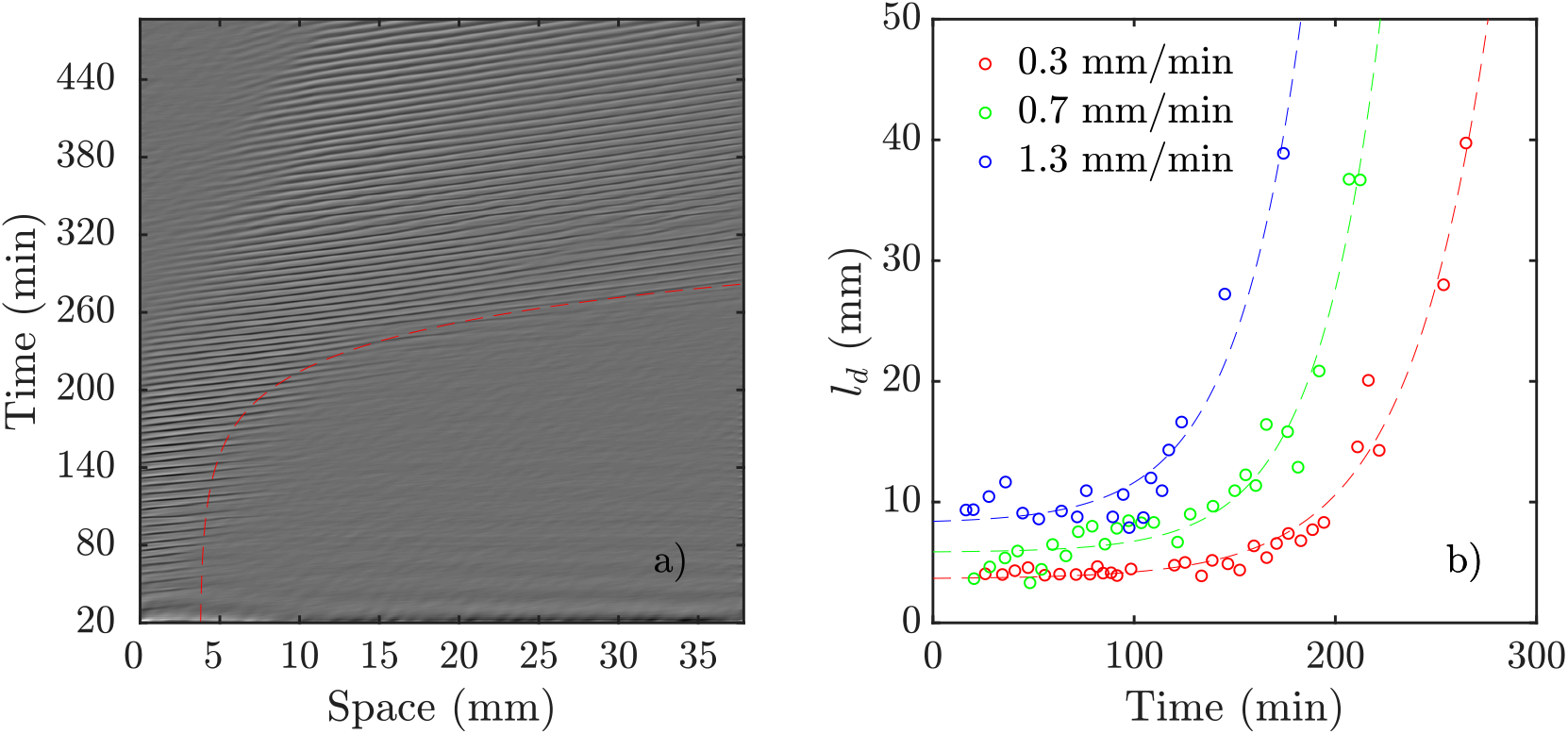
Pattern formation in experiments consisting of a channel filled with PdsA^-^ cells while the reservoir was filled with Ax2 cells starved for 4 hours (see movie S9). a) An exemplary space-time plot at *V_f_* = 0.7 mm/min with a wave period of 6.2 ± 0.3 min. The red dashed line illustrates the change of decay length over time (see movie S9). In b) the decay length is plotted vs experiment time for different flow speeds. Dashed lines show least-squares fit of exponential functions to the data of the same color. For each data set a function of the form of *f*(*x*) = *a* · exp(*b* · *x*) + *c* was used. The fitting values are: *a*_0.3_ = 0.0474 mm, *b*_0.3_ = 0.0250 min^-1^, *c*_0.3_ = 3.6374 mm, *a*_0.7_ = 0.0396 mm, *b*_0.7_ = 0.0316 min^-1^, *c*_0.7_ = 5.8376 mm and *a*_1.3_ = 0.1636 mm, *b*_1.3_ = 0.0303 min^-1^, *c*_1.3_ = 8.2406 mm.

### Numerical Simulations of PdsA^-^ Cells

We performed numerical simulations of the three-component MG model (Eq. 1) at different flow speeds. We selected *σ* and *k_e_* as control parameters since they account for the production and degradation of extracellular cAMP, respectively. Depending on these two parameters, this system can have stable, bistable and convectively unstable (CU) regimes, as is shown in the phase diagram in Fig. 11a. We fixed parameters *σ* and *k_e_* to be in the stable regime of the phase diagram where the boundary-driven oscillations exist. Note that BDOs exit in the CU regime too, where we have shown (1) that holding the upstream boundary to a zero concentration of cAMP produces an instability that sends periodic wave trains downstream. The wave generation mechanism works by first advecting the cAMP downstream, thus depleting the upstream area of cAMP. This low concentration of the chemoattractant destabilizes the cells close to the upstream boundary, which react by releasing a pulse of cAMP. For a careful analysis of BDOs in the CU regime, we refer the reader to Ref. (1).

**Figure 11:**
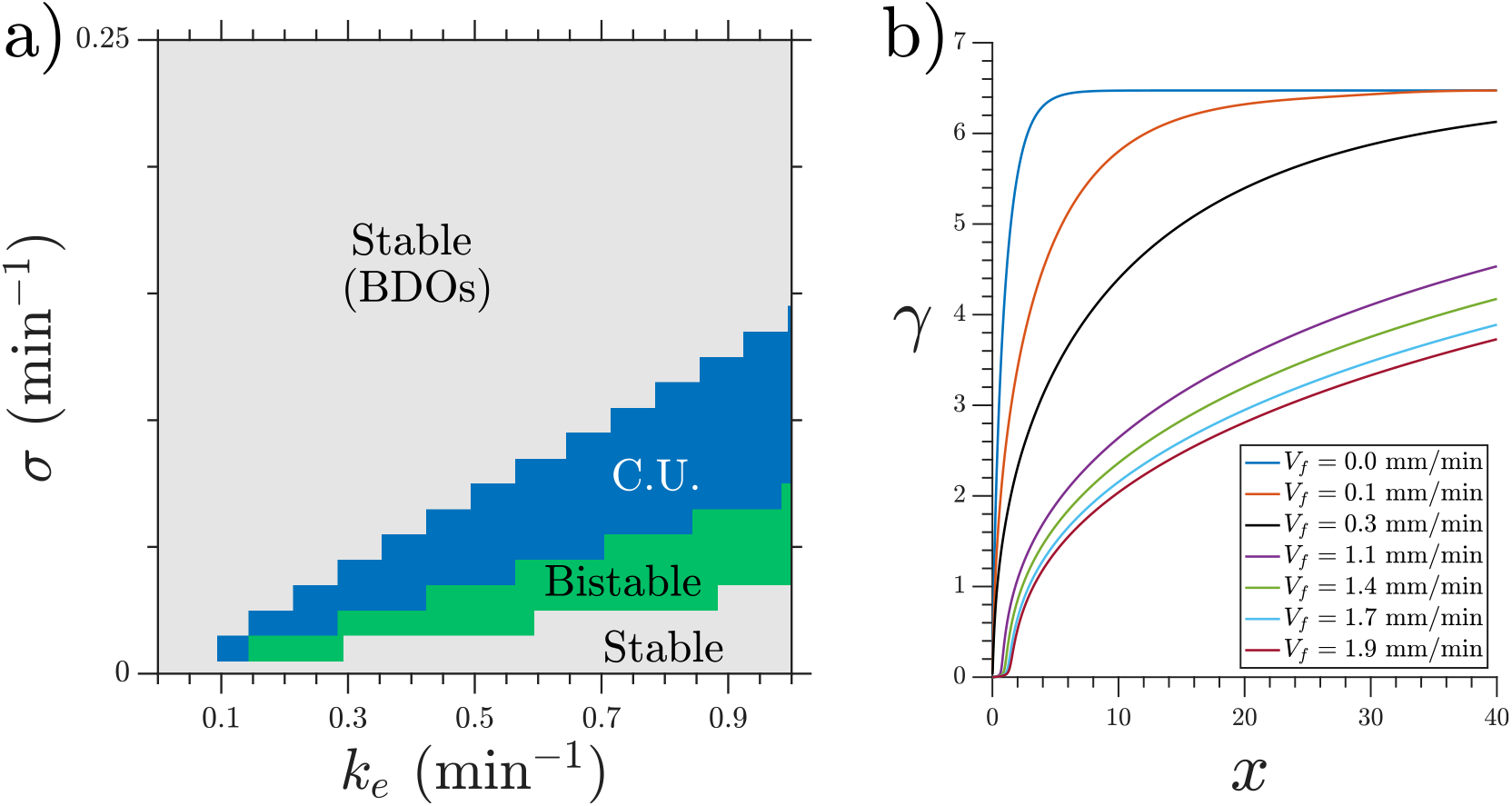
a) Phase diagram of the MG model showing the region at which BDOs exist. Note that BDOs exist in both stable and connectively unstable (CU) regimes. b) Steady state cAMP concentration *γ* obtained at *σ* = 0.3 min^−1^ and *k_e_* = 0.01 min^−1^. BDOs appear in the velocity range of 0.4 mm/min ≤ *V_f_* ≤ 0.8 mm/min.

To mimic our experimental setup, we performed numerical simulations in a channel filled with PdsA^-^ cells with imposition of a Dirichlet boundary upstream. To account for the low degradation rate of the cells, we consider small values of degradation rate *k_e_*. Numerical simulations showed that at low-speed flows the system reaches a time independent steady state. This solution is a monotonic curve that increases from a zero cAMP concentration at the channel upstream boundary up to a very high concentration in the channel downstream end (see Fig. 11b). At higher velocities of the advecting flow an instability appears, and the system’s solution is no longer time independent. The upstream boundary periodically emits waves that decay as they travel along the channel, similar to those observed in experiments. Examples of boundary-driven waves at three different flow speeds are shown in Fig. 12a-c and supplementary movies S10-S12. We emphasize that this instability only appears when an absorbing boundary condition in the upstream boundary is enforced, thus emulating the advection of cAMP-free buffer.

**Figure 12:**
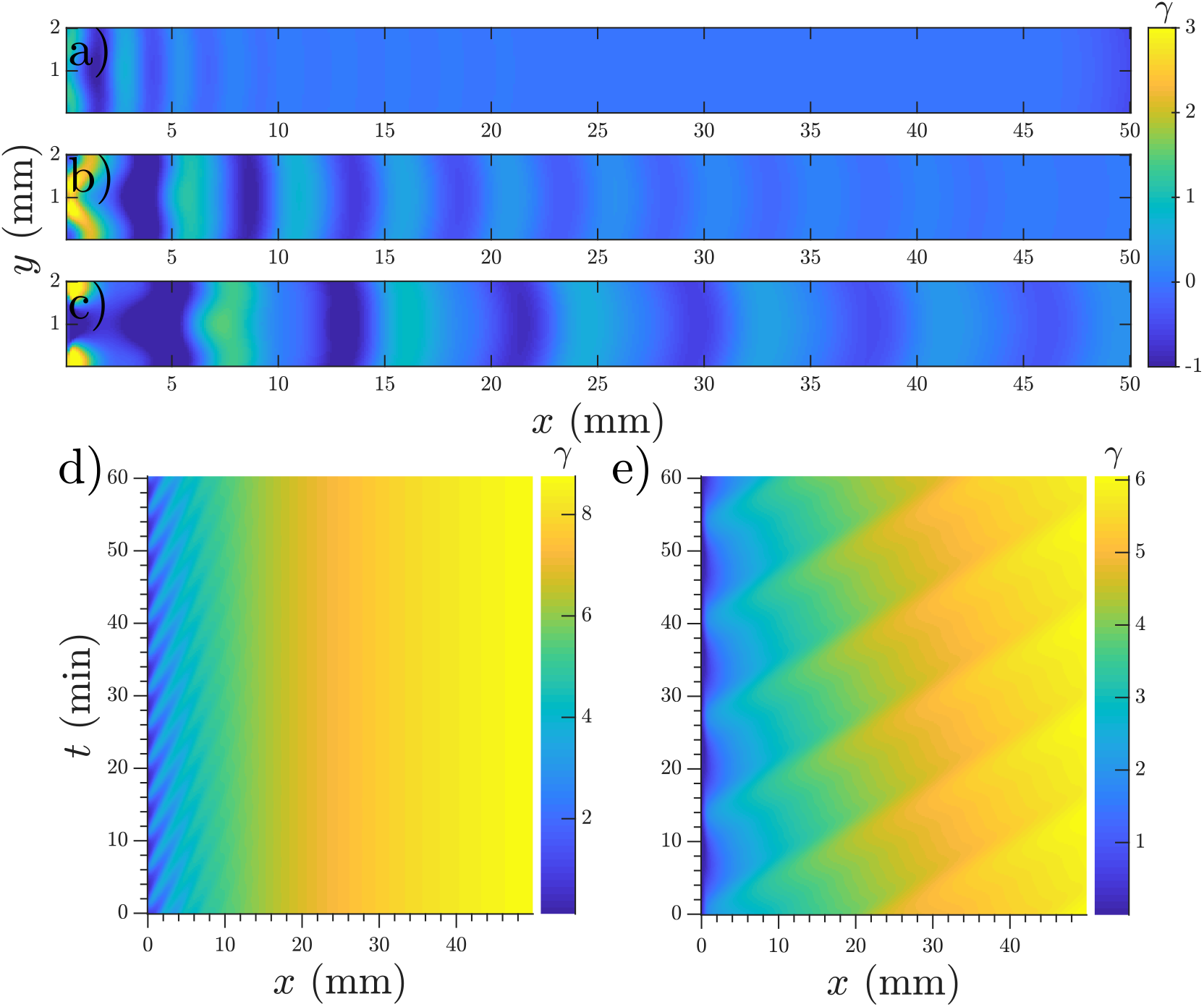
Boundary-driven waves developed in a channel with Dirichlet boundary condition at *x* = 0. a) *V_f_* = 0.5 mm/min, b) *V_f_* = 0.8 mm/min and c) *V_f_* = 1.1 mm/min (see movies S10-S12). Waves penetrate over longer distances at higher flow velocities. d-e) Space-time plot of boundary-driven waves at imposed flow velocities of 0.5 mm/min and 2 mm/min showing higher periods at higher flow speeds. Other parameters are *σ* = 0.6 min^−1^ and *k_e_* = 0.01 min^−1^. Note that in a)-c) the mean value of *γ* is subtracted to enhance the contrast of the waves. As a result, *γ* finds negative values and its range is different from space-time plots in parts d)-e).

We studied the properties of these waves in a 1D geometry along a range of cAMP production rate intensities (*σ*) and velocities, while keeping the degradation of cAMP very low (*k_e_* = 0.01 min^-1^). The properties of these waves are summarized in Fig. 13; it can be seen that the period of the advected waves increases with the advecting flows, while the characteristic length *l_d_* increases with flow velocity. The existence of these waves is restricted to a range of velocities, which increases to include faster flows at higher production rates of cAMP. Once the advecting flows are too fast for this mechanism to exist another numerical scheme needs to be used to periodically inject waves into the system. The high speed behavior is consistent with the previously reported behavior of WT (24) where the period is independent of the advecting flows with *T* ≈ 5 – 6 min.

**Figure 13:**
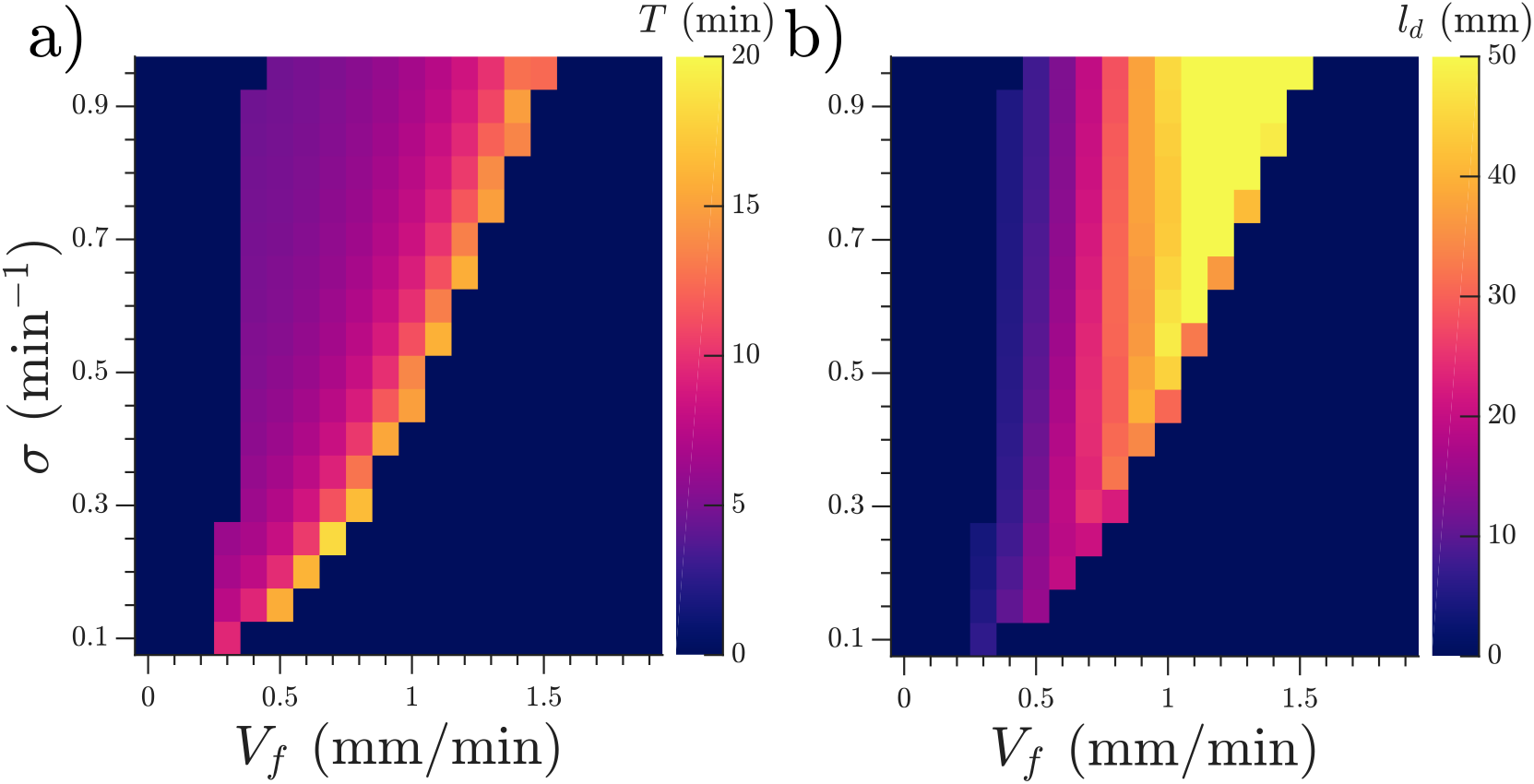
a) Period *T* and b) characteristic decay length *l_d_* of the observed waves as functions of the velocity of the advecting flow *V_f_* and the production rate *σ* of cAMP. Note that simulations are performed in a 1D geometry. Degradation rate of cAMP is *k_e_* =0.01 min^-1^ and channel length is *L* = 50 mm.

We also performed numerical simulations emulating the experiments with WT cells in the inlet. In those simulations the parameters were fixed as *k_e_* = 5.0 min^-1^ and *σ* = 0.55 min^-1^ for *x* < *l* with *l* = 1 mm to represent the inlet. Under these parameters, the inlet oscillates with a fixed period and emits waves to the rest of the channel. In simulations including phosphodiesterase advection and WT cells in the inlet we observed that the secreted phosphodiesterase was advected quickly downstream and therefore the phosphodiesterase concentration could be assumed constant along the channel during most of the experiment. Given this phosphodiesterase distribution, in the experiments when the cells are not prestarved and signaling starts after 3 hours the system behaves like a channel filled with WT.

In the case where the WT cells are prestarved and start to fire immediately after being placed in the channel we found that an increasing production rate of cAMP for the PdsA^-^ cells reproduced the increasing decaying length in the system. Simulations for different production rates with fixed degradation rates (fixed at WT values *k_e_* = 5.0 min) were performed, and their decay length increased with increasing production rate, as is shown in Fig. 14a. An example of a simulation where the production rate increased over time as *σ* = *Kt* (*K* is a constant) can be seen in Fig. 14b; there, the decay length increases over time (see movie S11).

**Figure 14:**
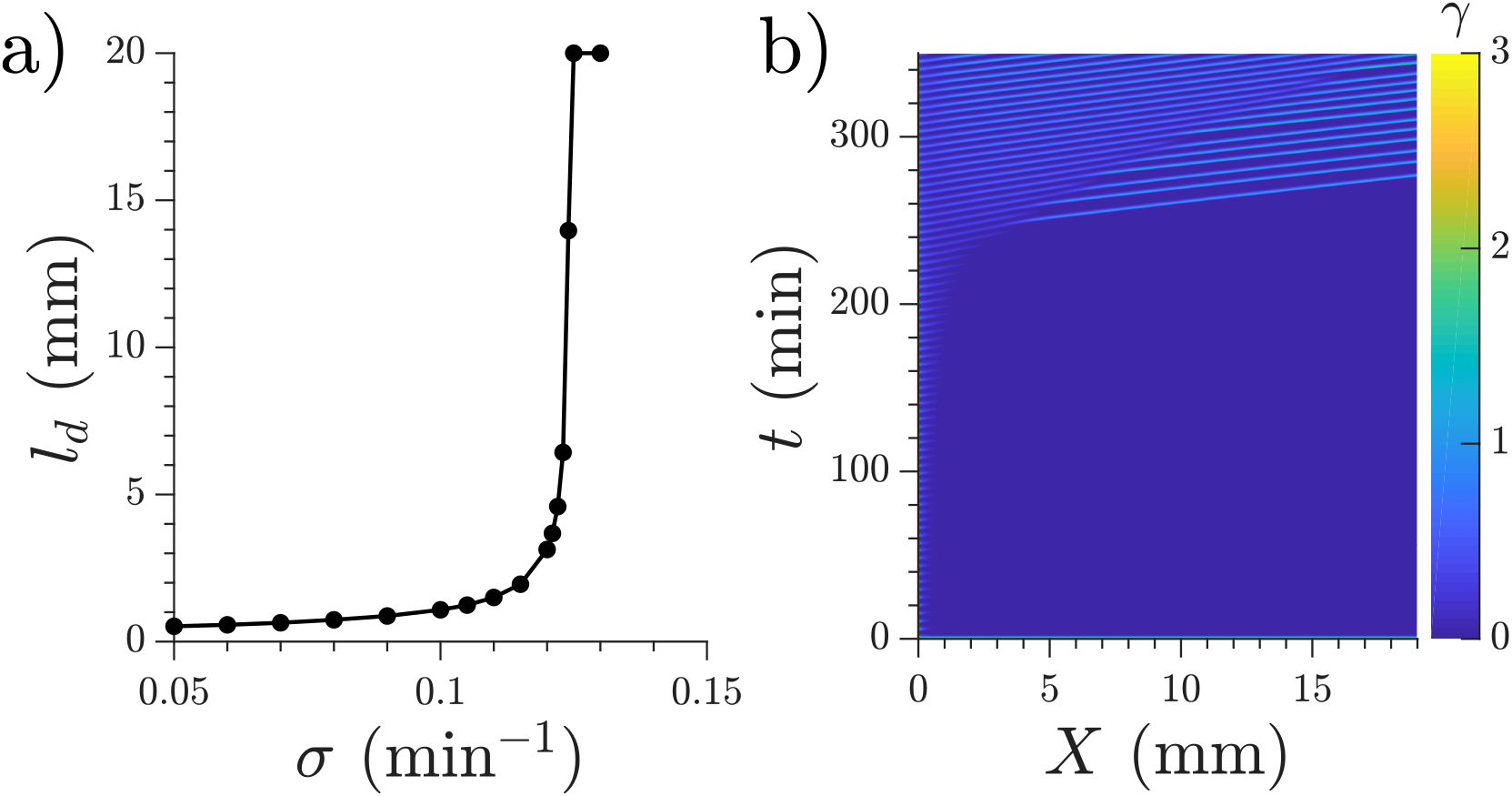
a) Characteristic length *l_d_* for the waves produced at the inlet by the WT cells. The degradation rate given by the effect of phosphodiesterase is fixed at *k_e_* = 5.0 min^-1^ for the whole channel. b) Space-time representation of a numerical simulation with WT cells in the inlet (not shown) and linearly increasing cAMP production rate in the rest of the channel, *σ* = 0.001 · *t*. Flow velocity is 0.7 mm/min (see movie S13).

## DISCUSSION

We have analyzed and characterized the properties of boundary-driven waves in a population of PdsA deficient cells. Our experiments show that an influx of cAMP-free buffer does not only recover signaling in PdsA^-^ cells, but also that these cells are capable of signaling and aggregation once the flow has been switched off. In our flow-through microfluidic setup filled with PdsA^-^ cells we observed cAMP waves after 3 hours of starvation with flow, independent of whether or not they were previously starved in a shaker. This confirms that reduced levels of cAMP are necessary for their development. These waves have a finite decay length inside the channel which grows with the imposed flow velocity. Boundary-driven waves further develop the cells, and thereby waves reach longer distances along the channel after around 6.5 hours throughout the experiment. Since cells continue signaling after the flow is stopped, it is plausible that other types of phosphodiesterases such as PDE4 take over the role of PdsA during aggregation (12), having a higher activity than in the normal presence of PdsA. Other biochemical experiments are necessary to confirm this hypothesis. Interestingly, the period of the spontaneous waves observed in the recovered population of PdsA^-^ cells was around 15-25 min which is much higher than the natural period of WT cells (~ 6 min). The rescued cells were located on the upstream of the channel, consistent with the area covered by the decaying boundary-driven waves. In addition to the penetration length of the waves, the wave amplitude was also a determining factor in triggering cell aggregation as the cells located downstream experiencing low amplitude waves fail to aggregate.

We performed numerical simulations using the reaction-diffusion model proposed by Martiel and Goldbeter (22). Our simulations in the stable regime of the phase diagram confirms the existence of a boundary-driven instability that produces waves in this system. A flow of cAMP-free buffer provides the destabilizing mechanism necessary for wave production, thus showing boundary-driven oscillations. At small flow speeds the clean buffer produces depletion of cAMP in the upstream edge of the channel, thus allowing the cells located there to fire and produce a cAMP wave. As this wave travels downstream the basal concentration of cAMP increases and the wave loses amplitude, decaying as it travels. At higher flow speeds more clean buffer penetrates the system, thus the basal cAMP concentration increases more slowly along the channel (smaller gradients), therefore allowing the waves to travel further. We systematically measured the penetration length of the waves and found that it increases with the imposed flow velocity, which is in agreement with our experiments. In our simulations there is a maximum velocity such that the boundary-driven oscillations exist. This is in disagreement with our experimental observations, where at higher flow speeds wave generation continues to occur. We believe that the range of existence of this instability is marked by the change in the oscillating period observed in experiments (see the vertical line in Fig. 2c). For *V_f_* < 0.6 mm/min the wave period is large and changes with the advecting flow, while for *V_f_* > 0.6 mm/min it is constant around *T* ≈ 5 – 6 min, therefore these BDOs exist for *V_f_* < 0.6 mm/min. To test this hypothesis we performed numerical simulations including a constant firing of the cells with *T* = 6 min. For the range of velocities where the boundary-driven oscillations exist, the period of these oscillations prevailed and the firing had no effect on the system. For higher velocities the constant firing set the system’s period.

Moreover, we used WT cells in the upstream end of the channel to rescue PdsA^-^ cells. We observed waves with increasing penetration length that propagated inside the channel. This behavior was reproduced in our simulations with WT cells at the reservoir with an increasing cAMP production rate of PdsA^-^ cells everywhere in the channel. Therefore, we attribute this increasing penetration length to the gradual development of the PdsA^-^ cells in the channel. In our experiments, if the WT cells at the reservoir are not prestarved, then the WT cells develop along with the PdsA^-^ cells and the flow-driven waves appear after 3 hours and are capable of penetrating throughout the channel.

In summary, we have shown experimentally and by means of numerical simulations that by imposition of Dirichlet boundary condition in a reaction-diffusion-advection system, an instability occurs which generates periodic waves traveling downstream. We emphasize that the observed phenomena is different from periodic wave trains emitted by Dirichlet boundary condition in oscillatory reaction-diffusion systems (29), in that the waves are induced in a stable reaction-diffusion system and the wave amplitude decays as the waves propagate downstream. We expect this type of boundary-driven instability to be a generic mechanism to produce continuous periodic influx of wave trains in an otherwise stable reaction-diffusion system.

## AUTHOR CONTRIBUTIONS

A.G. designed the research. E.V.H. designed and carried out numerical simulations. T.E. performed experiments. E.V.H., T.E. and A. G. analyzed data. A.G., E.V.H. and T.E. wrote the manuscript. E.V.H. and T.E. contributed equally.

## ACKNOWLEDGMENTS

The authors thank V. Zykov and J. McEnnay for a careful reading of the manuscript and M. S. Müller, S. Romanowsky and K. Gunkel for their cheerful help with the preparation of the cells. T. E. acknowledges Deutsche Forschungsgemeinschaft (DFG), project number GH 184/1-1. E.V.H. thanks the Deutsche Akademische Austauschdienst (DAAD), Research Grants—Doctoral Programs in Germany. A.G. acknowledges MaxSynBio Consortium, which is jointly funded by the Federal Ministry of Education and Research of Germany and the Max Planck Society.

